# Predictive model in the presence of missing data: the centroid criterion for variable selection

**DOI:** 10.1101/420943

**Authors:** Jean Gaudart, Pascal Adalian, George Leonetti

## Abstract

**Introduction:** In many studies, covariates are not always fully observed because of missing data process. Usually, subjects with missing data are excluded from the analysis but the number of covariates can be greater than the size of the sample when the number of removed subjects is high. Subjective selection or imputation procedures are used but this leads to biased or powerless models.

The aim of our study was to develop a method based on the selection of the nearest covariate to the centroid of a homogeneous cluster of covariates. We applied this method to a forensic medicine data set to estimate the age of aborted fetuses.

**Analysis:** *Methods:* We measured 46 biometric covariates on 50 aborted fetuses. But the covariates were complete for only 18 fetuses. First, to obtain homogeneous clusters of covariates we used a hierarchical cluster analysis. Second, for each obtained cluster we selected the nearest covariate to the centroid of the cluster, maximizing the sum of correlations 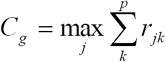 (the centroid criterion). Third, with the covariate selected this way, the sample size was sufficient to compute a classical linear regression model. We have shown the almost sure convergence of the centroid criterion and simulations were performed to build its empirical distribution. We compared our method to a subjective deletion method, two simple imputation methods and to the multiple imputation method.

*Results:* The hierarchical cluster analysis built 2 clusters of covariates and 6 remaining covariates. After the selection of the nearest covariate to the centroid of each cluster, we computed a stepwise linear regression model. The model was adequate (R^2^=90.02%) and the cross-validation showed low prediction errors (2.23 10^−3^). The empirical distribution of the criterion provided empirical mean (31.91) and median (32.07) close to the theoretical value (32.03). The comparisons showed that deletion and simple imputation methods provided models of inferior quality than the multiple imputation method and the centroid method.

*Conclusion:* When the number of continuous covariates is greater than the sample size because of missing process, the usual procedures are biased. Our selection procedure based on the centroid criterion is a valid alternative to compose a set of predictors.

## Introduction

Predictive models are widely used in life sciences. For clinical practice, usual statistical models like regression models are often build using observed covariates to estimate the value of a dependant variable. Particularly in anthropology and forensic sciences, predictive models are mostly used to predict age and gender of subjects from different biometrics covariates. However, all covariates are not always fully observed since data measurements can be difficult (for example in case of superposition in radiographs) and because forensic specialists frequently have to deal with incomplete human remains. Usually, subjects with missing data are excluded from the analysis (Complete-Case Analysis). But first, if the missing data depend on a non-random process related to the dependant variable, excluding this selective group leads to biased models [**1**]. Second, this practice reduces the statistical power of the analysis. Third, in some situations the number of removed subjects is unacceptably high: even if the missing rates per covariate are low, few subjects may have complete data for all covariates leading to analyze a number of covariates greater than the size of the sample with complete data. Then, statistical analysis and result interpretation are tricky: usual statistical methods like stepwise regression methods are not able to select the predictive covariates. To compose a set of predictors in this context, covariates are often selected via ad hoc and subjective means mostly based on the number of available data for each covariate leading to biased models.

To avoid this problem imputation methods consist on substituting the missing data with plausible values. The most popular simple imputation practice is the unconditional mean substitutions where missing data are replaced by the mean 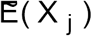 estimated from the observed values of each covariate ***X*_*j*_** [**2, 3**]. This method preserves the observed sample means but under-estimates variances and covariances. Another simple imputation method substitutes each missing data by conditional means estimated from regression models (e.g. linear regression) where the covariate ***X*_*j*_** is estimated using the dependant variable 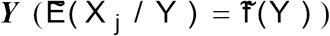 or other0020covariates 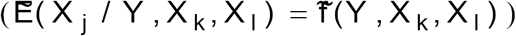. But this practice biases again the models over-estimating the correlations. Another method called multiple imputations has been used in several applications [**4, 5, 1, 6, 7, 8**]. This technique generates imputations using Expectation-Maximization (EM) algorithm or Data Augmentation (DA) algorithm [**9**]. The results are combined taking into account the imputation uncertainty. However, missing data are generated from models based on the observed data and therefore multiple imputation increases colinearity [**3**].

The aim of this study was to develop, in this context, a method to select covariates using neither imputations nor subjective procedures and reducing amounts of information to be discarded. This method is based on the selection of the nearest covariate to the centroid of a homogeneous cluster of covariates. This selected covariate minimizes the distance to the other covariates and therefore a large amount of the cluster information is preserved. We applied this method to a forensic medicine data set in order to estimate the age of aborted fetuses.

## Analysis

### 1. Material and methods

In case of abortions caused by traumatic situations, the fetal age has to be accurately determined. Indeed, in the french law the judgment and the prejudice appreciation differ in step with the fetal age. Fetal age can be determined using thigh-bone lengths [**10**] or foot lengths [**11**] as well as cranium bones measures. For the latter procedure of determination, 46 biometric covariates were measured (cranium bone lengths or angular measures [**12**, **13**]) on 50 aborted fetuses. But, because of the traumatic situations, it was not possible to measure all the covariates on the whole fetus sample. The measures were complete for only 18 fetuses.

First, to obtain homogeneous clusters of covariates we used a hierarchical cluster analysis using Pearson correlations as similarity measures.

Second, for each obtained cluster of covariates we selected the nearest covariate to the centroid of the cluster. We have determined that the nearest covariate to the centroid is the covariate which maximizes the sum of correlations. Indeed [appendix 1], the squared distance between a covariate and the centroid is given by the Torgerson formula [**14**, **15**]:

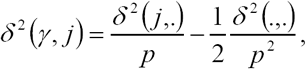

Where 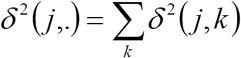 and 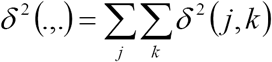 and *p* is the number of covariates.

If *δ* ^2^ (*j, k*) = 2(1– *ρ* _*jk*_) is the squared distance between two covariates, where *ρ*_*jk*_ is the correlation coefficient between two covariates,

Then 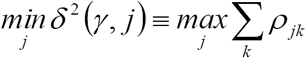

Furthermore, we have shown that this criterion 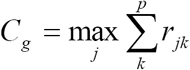 converges almost surely [appendix 2].

Third, with the covariate selected this way, the sample size was sufficient to compute a stepwise linear regression model. The results were analyzed by cross-validation procedure and goodness-of-fit was estimated by the rate of explained variability R^²^.

In the presented statistical context (finite sample size and *ρ*_*jk*_≠0), the distribution of the empirical correlation coefficient, *r*_*jk*_, is not simply usable [**16**, **17**]. Then, it is not possible to formalize the distribution of the maximum of the empirical correlation sum 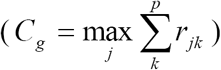. Therefore we studied the empirical distribution of *C*_*g*_ by simulations. Using the variance-covariance matrix of a cluster of *p* covariates issued from our sample, we simulated the same number *p* of covariates. Those covariates were extremely correlated and were normally distributed. To determine the number of simulations, we used, for each centile, the relative errors between a centile *c*(*i*; *n*) issued from the empirical distribution simulated with *n* simulations, and the same centile *c*(*i*; *n* –100) issued from the empirical distribution simulated with *n-100* simulations:

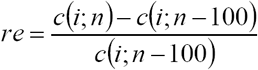

The simulation was stopped when *re* ≤ 0.2% for each centile, and the empirical distribution was considered as stable.

We compared the resulting model with 4 other methods:

i. The deletion procedure deleting covariates until the sample size is greater than the number of covariates;
ii. The simple unconditional mean imputation method, imputing the estimated mean of each covariate to missing data (for each covariate **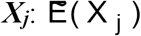**);
iii. The simple conditional mean imputation method, imputing to missing data the predicted value resulting from a simple linear regression on the variable Age (for each covariate 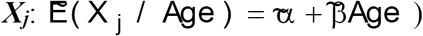;
iv. The multiple imputation method: to generate imputations we used the DA algorithm (broadly described by **9**). For this purpose, Schaffer’s Norm^®^ freeware was employed. Missing values were replaced by 20 simulated values in order to assure the efficiency of the procedure [**2**]. The 20 sets of imputations reflected the uncertainty about the true values of missing data. Each of the 20 completed data sets was analyzed using standard linear regression method. The results were combined into a single inference according to the procedure described for example in [9] [2] or [**4**].

The datasets, obtained by these four methods, were analyzed by stepwise linear regression. The goodness-of-fits of the resulting models were compared using the rates of explained variability and the accuracies were compared by the prediction errors provided by cross-validations.

### 2. Results

#### 2.1 Centroid method of selection

The hierarchical cluster analysis (fig. 1), using 46 covariates, built 2 clusters with respectively 36 and 4 clustered covariates. The 6 remaining covariates (AFW: Weisbach angle, APN: nasal angle, ATO: foramen magnum angle, ASP: nasion-sphenion-basion angle, NKB: nasion-klition-basion angle, ABO: angle formed by the straight line nasion-opisthion and the Francfort section) appeared to be separate from the other covariates. In the first cluster (36 covariates) the *C*_*g*_ criterion allowed us to select the covariate called LGB (glabella-basion length). That is, this covariate maximized the correlation sum and therefore was the nearest covariate to the centroid of the first cluster. The covariate LGB (fig. 2) represents the length between the frontal bone and the foramen magnum. This length is a usual and standardized measure [**18**] and, for fetuses, LGB can be easily measured by scanner. In the same way, the *C*_*g*_ criterion allowed us to select the covariate called APA (alveolar profile angle [**13**, **18**]) in the second cluster composed of 4 covariates.

On the whole we obtained 8 covariates with a sample size of 35 fetuses (free of missing data) to compute a stepwise linear regression model. We obtained the following model (fig. 3):

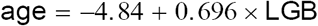

(age in weeks of amenorrhea, LGB in millimeter).

**Figure 1:**
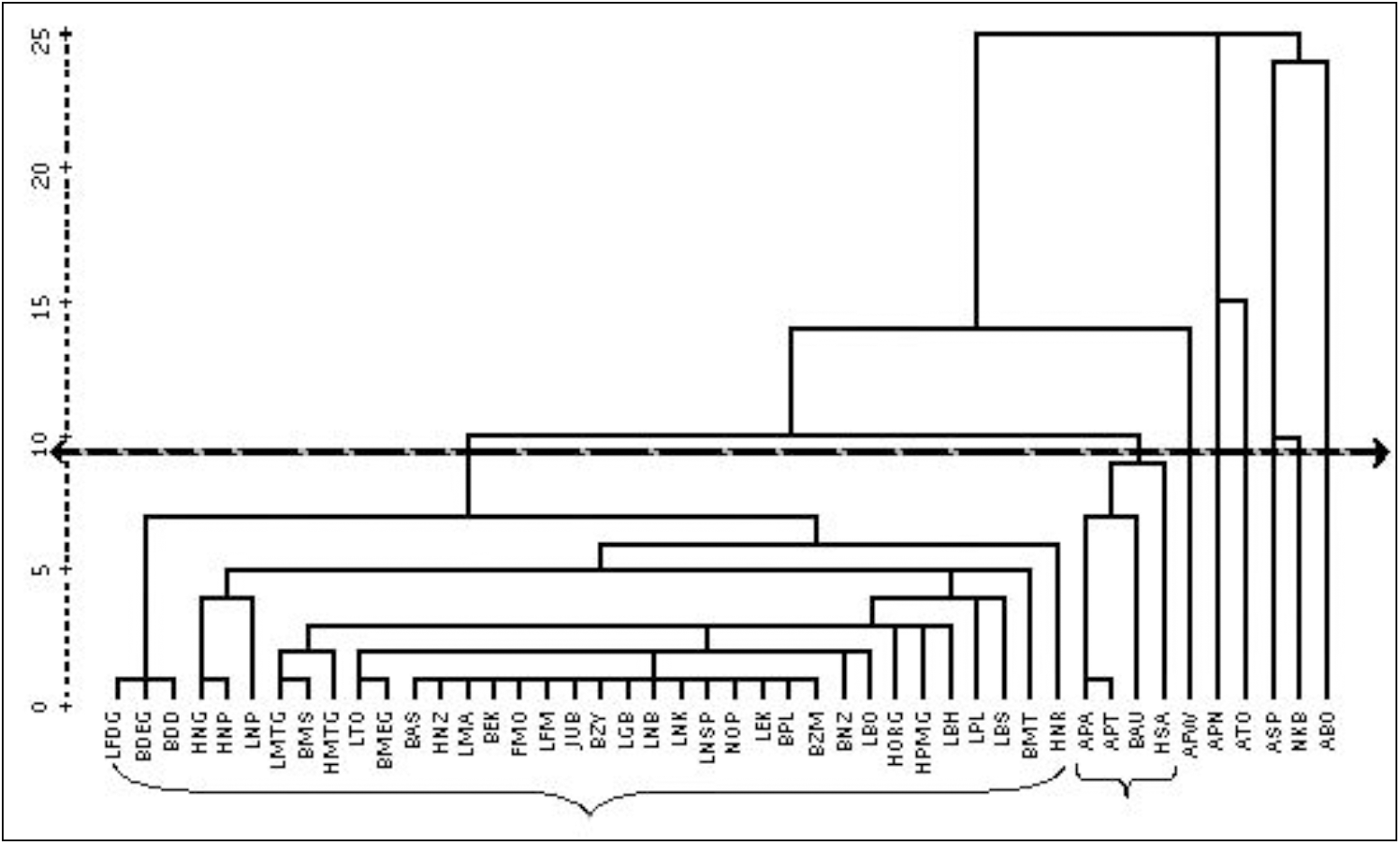
hierarchical cluster analysis using the correlation coefficient of Pearson as similarity measure. We obtained 2 homogeneous clusters of covariates and 6 remaining covariates.

**Figure 2:**
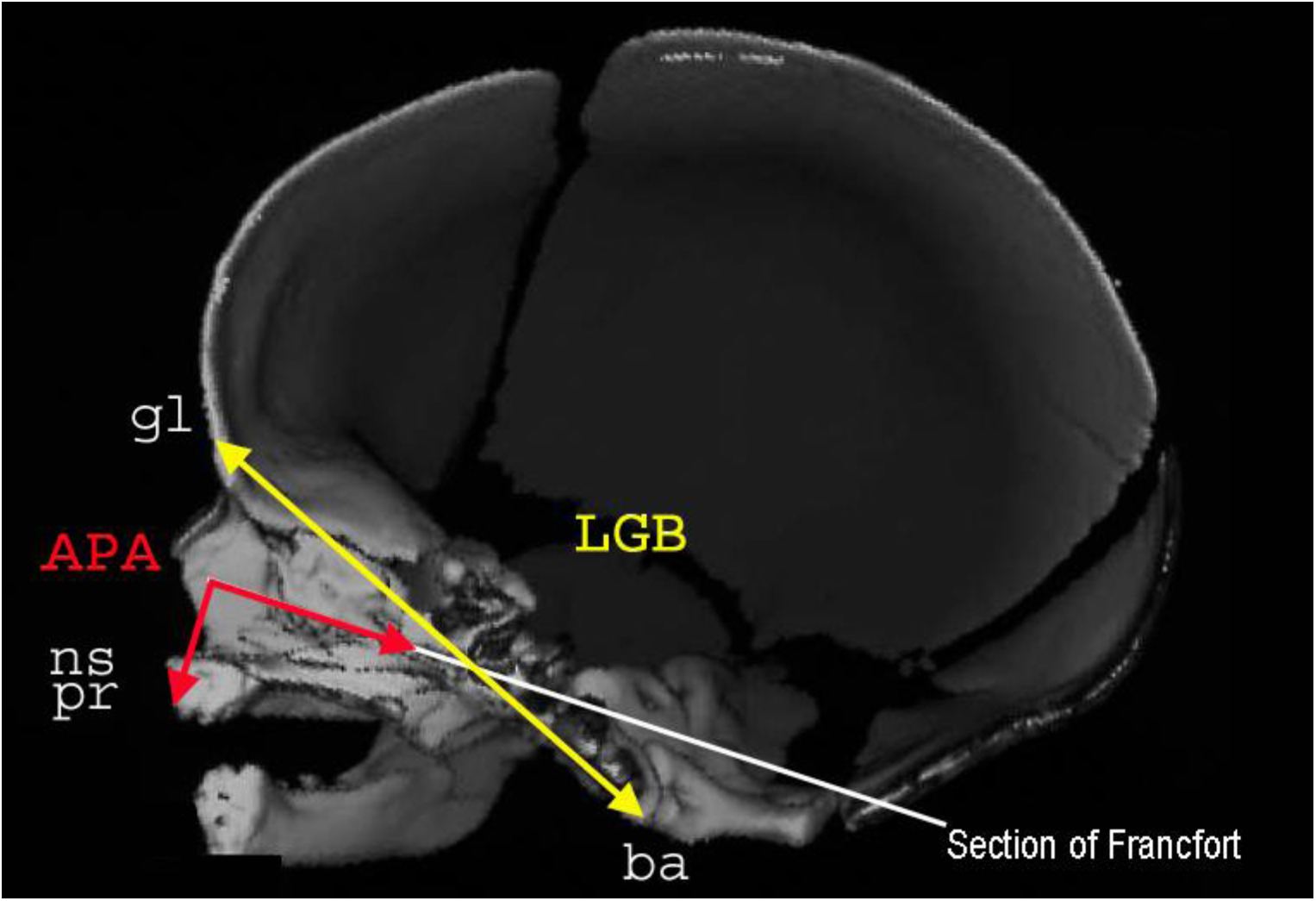
scanner of a male fetus of 37 weeks of amenorrhea: sagittal section. *LGB*: glabella basion length *APA*: alveolar profile angle *gl*: glabella: most anterior midline point of the frontal bone, usually above the frontonasal suture. *ns*: nasospinal: intersection between the midline and the tangent to the margin of the inferior nasal aperture, at the lowest points. *pr*: prosthion: midline point of the most anterior point of the maxillae alveolar process. *ba*: basion: midline point of the anterior margin of the foramen magnum.

**Figure 3:**
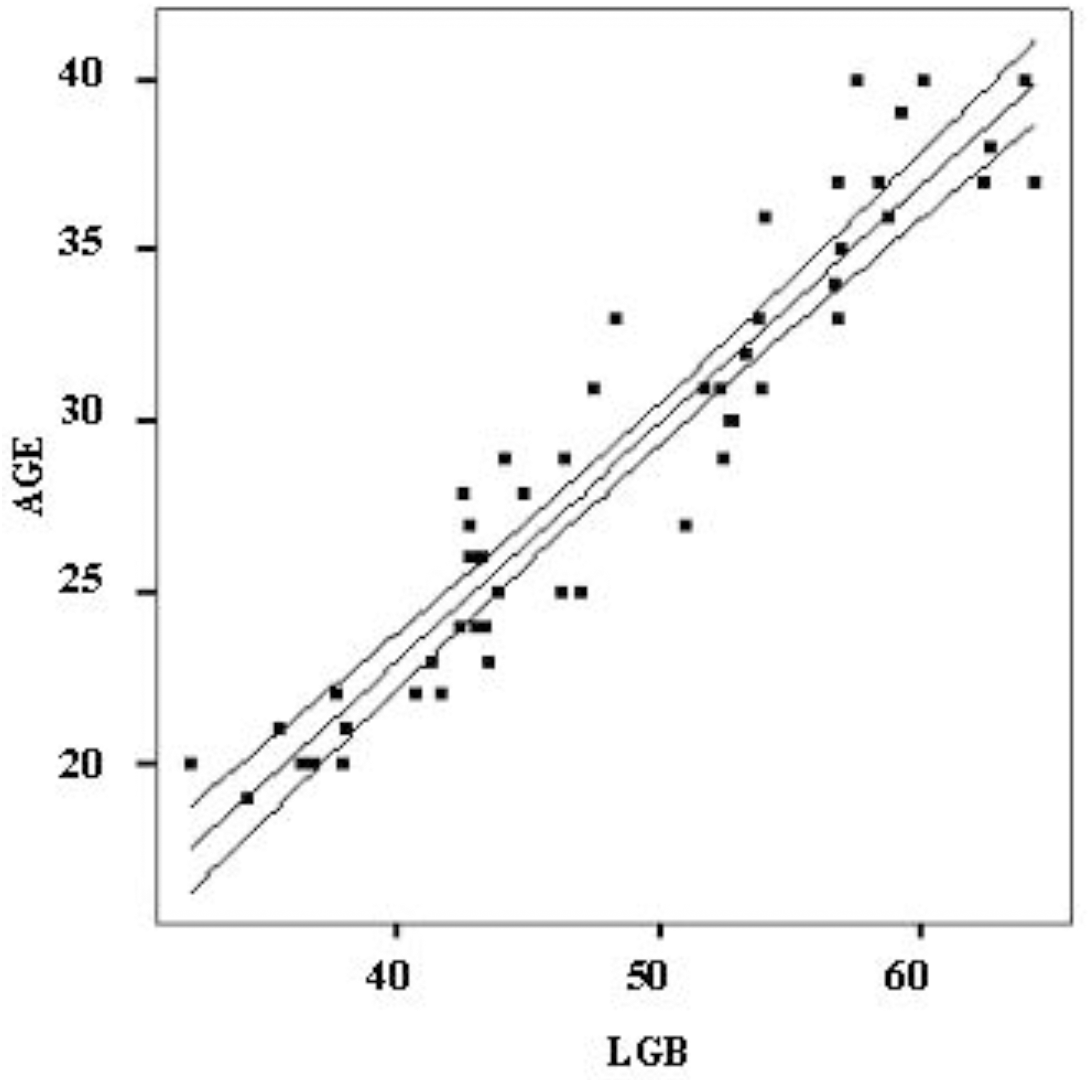
scatter plot of the age (in weeks of amenorrhea) and the LGB (in millimeter). The regression model is shown with its 95% confidence interval.

The model was adequate (*R*^*2*^=90.02%) and the cross-validation showed very low prediction errors (mean: 2.23 10^−3^ CI95%[−0.58; 0.58]) (table 1).

**Table 1:**
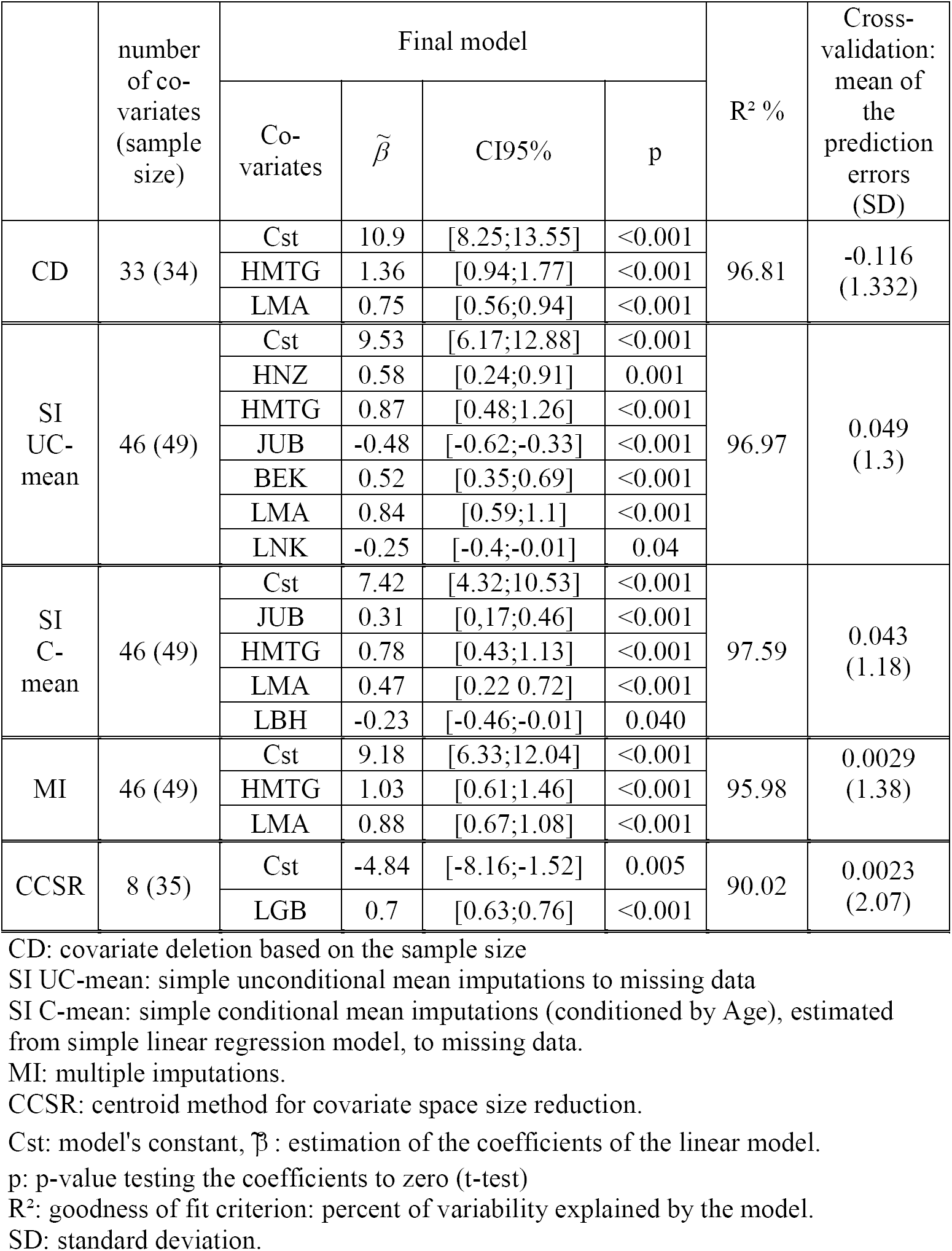
Results of the four modeling procedures.

|  | number of co-<br>variates<br>(sample size) | Final model |  |  |  | R<sup>2</sup> % | Cross-validation:<br>mean of the<br>prediction errors<br>(SD) |
| --- | --- | --- | --- | --- | --- | --- | --- |
| | | Co-<br>variates | $\tilde{\beta}$ | CI95% | p | | |
| CD | 33 (34) | Cst | 10.9 | [8.25;13.55] | <0.001 | 96.81 | -0.116<br>(1.332) |
|  |  | HMTG | 1.36 | [0.94;1.77] | <0.001 |  |  |
|  |  | LMA | 0.75 | [0.56;0.94] | <0.001 |  |  |
| SI<br>UC-mean | 46 (49) | Cst | 9.53 | [6.17;12.88] | <0.001 | 96.97 | 0.049<br>(1.3) |
|  |  | HNZ | 0.58 | [0.24;0.91] | 0.001 |  |  |
|  |  | HMTG | 0.87 | [0.48;1.26] | <0.001 |  |  |
|  |  | JUB | -0.48 | [-0.62;-0.33] | <0.001 |  |  |
|  |  | BEK | 0.52 | [0.35;0.69] | <0.001 |  |  |
|  |  | LMA | 0.84 | [0.59;1.1] | <0.001 |  |  |
|  |  | LNK | -0.25 | [-0.4;-0.01] | 0.04 |  |  |
| SI<br>C-mean | 46 (49) | Cst | 7.42 | [4.32;10.53] | <0.001 | 97.59 | 0.043<br>(1.18) |
|  |  | JUB | 0.31 | [0.17;0.46] | <0.001 |  |  |
|  |  | HMTG | 0.78 | [0.43;1.13] | <0.001 |  |  |
|  |  | LMA | 0.47 | [0.22 0.72] | <0.001 |  |  |
|  |  | LBH | -0.23 | [-0.46;-0.01] | 0.040 |  |  |
| MI | 46 (49) | Cst | 9.18 | [6.33;12.04] | <0.001 | 95.98 | 0.0029<br>(1.38) |
|  |  | HMTG | 1.03 | [0.61;1.46] | <0.001 |  |  |
|  |  | LMA | 0.88 | [0.67;1.08] | <0.001 |  |  |
| CCSR | 8 (35) | Cst | -4.84 | [-8.16;-1.52] | 0.005 | 90.02 | 0.0023<br>(2.07) |
|  |  | LGB | 0.7 | [0.63;0.76] | <0.001 |  |  |

CD: covariate deletion based on the sample size
SI UC-mean: simple unconditional mean imputations to missing data
SI C-mean: simple conditional mean imputations (conditioned by Age), estimated from simple linear regression model, to missing data.
MI: multiple imputations.
CCSR: centroid method for covariate space size reduction.
Cst: model's constant, $\tilde{\beta}$ : estimation of the coefficients of the linear model.
p: p-value testing the coefficients to zero (t-test)
R<sup>2</sup>: goodness of fit criterion: percent of variability explained by the model.
SD: standard deviation.

#### 2.2 Simulations

Using the variance-covariance matrix of the first homogeneous cluster of covariates, we simulated 36 correlated and normally distributed covariates.

With 1000 simulations (fig. 4) the empirical distribution of *C*_*g*_ was stable that is the relative errors of each centile were lower than 0.02. The empirical mean (31.91 CI95% [31.86; 31.96]) and the empirical median (32.07) were close to the theoretical value (32.03) in spite of the small size of the simulated samples (*n*=18 subjects). The empirical variance (0.56) and the empirical range (minimum=26.75; maximum=33.22) were also small.

**Figure 4:**
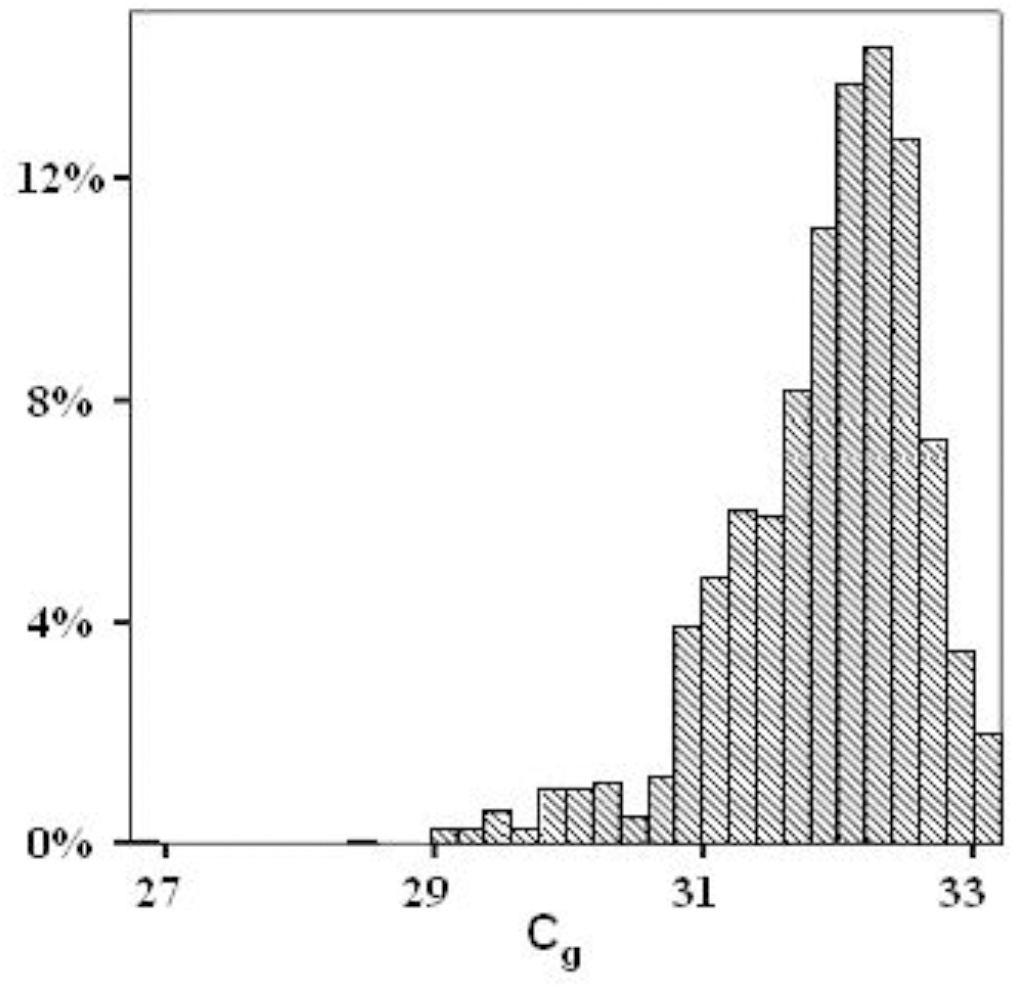
empirical distribution of 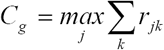, 1000 simulations of 36 correlated and normal distributed covariates.

#### 2.3 Comparisons

The four other methods provided four different groups of selected covariates. These groups were analyzed using stepwise linear regression procedure.

i. The first method provided covariate deletions based on the sample size: covariates composed of the larger number of missing data were deleted until the size of the sample (with complete data) was greater than the number of covariates. We obtained 33 fully observed covariates for 34 subjects. Stepwise linear regression provided a linear model composed of 2 covariates (HMTG: mastoid height and LMA: maxillo-alveolar length) (table 1). The rate of explained variability R^²^ was important (96.81%) but cross validation provided non-negligible prediction errors (mean:-0.116 CI95%[− 0.564;0.332]).
ii. The analysis of the sample completed with simple imputations of unconditional means provided a linear model composed of 6 covariates (HNZ: nasal height, HMTG, JUB: bijugal breadth, BEK: biorbital breadth, LMA, LNK: nasion-klition length). The R^²^-value was important 96.97% and cross validation provided low prediction errors (mean: 0.049 CI95%[−0.32;0.41]) (table 1).
iii. With the simple imputation method imputing conditional means (estimated from simple linear regression models) to missing data, stepwise linear regression procedure provided a final predictive model composed of 4 covariates (JUB, HMTG, LMA, LBH: basion-hormion length). The goodness-of-fit criterion R^²^ was high (97.59%) and cross validation provided low prediction errors (mean: 0.043 CI95%[−0.29;0.37]) (table 1).
iv. Finally, the multiple imputation method provided a model composed of 2 covariates (HMTG, LMA). The R^²^-value (95.98%) was slightly lower than the R²- values of the latter models but the cross validation provided very low prediction errors (mean: 0.0029 CI95% [−0.38;0.39]).

## 3. Discussion

When the number of continuous covariates is greater than the sample size, because of missing data process, imputations methods or ad hoc selections based on available data are commonly applied. Our study showed that in this context the selection method based on the centroid criterion is a valid alternative method. Indeed, the choice of the nearest covariate to the centroid of a cluster of covariates is a simple selection method which avoids resorting to imputations or ad hoc selections.

Complete Case Analysis cannot be applied and the covariate selection based on the sample size is not accurate as this method leads to biased models with high prediction errors. Simple imputation methods involve well-known biases [**1, 2, 3, 9**]. Unconditional mean imputations provide underestimated variances and covariances and introduce a conservative bias reducing the strength of the relationship between the dependant variable ***Y*** and the covariates ***X*_*j*_** completed this way. Conditional mean imputations provide overestimated correlations and over-fitted models. This method introduces costly bias increasing the strength of the relationship between the completed covariates ***X*_*j*_** and the dependant variable ***Y***. On the other hand, the multiple imputation procedure provided here the best model according to goodness-of-fit and prediction errors. Performing 20 imputations lead to an efficiency of 96.9% in the presence of more than 50% of missing data [**2**]. But MI is known to increase colinearity [**1**, **2, 3**, **4, 9, 19**]. It introduces a costly bias increasing the strength of the relationship between the dependant variable ***Y*** and the completed predictive covariates ***X*_*j*_**. Furthermore, different MI procedures can lead to different inferences [**7**]. Even if multiple imputation method does not much disturb the results of an explicative model (i.e. the presence or absence of relationship between the dependant variable and the covariates) imputations do modify predictions [**6**]. The selection method based on the centroid criterion is a good alternative finding an accurate model: even if the results displayed the lowest goodness-of-fit, the latter remained high (R²=90.02%) and cross validation provided the lowest prediction errors mean: 0.0023 CI95% [−0.58;0.58]). Missing data do not disturb our selection method as far as they are missing at random. The correlation was estimated between two covariates and for these covariates the number of data has to be sufficient to ensure the convergence of the correlation estimation.

We have proven the almost sure convergence of the centroid criterion [appendix 2]. But the speed of convergence depend on the sample size *n* and as well as the number *p* of covariates. If *p* is greater than *n*, even if the convergence of the two-by-two correlation coefficient is reached, the correlation matrix will be formally degraded. In a practical way, if *n* is sufficient, the convergence default will be weak.

For detecting homogeneous clusters of covariates we have chosen hierarchical cluster analysis. But other clustering procedures can be used, such as K-means clustering. In our study, in spite of small changes in the clusters of covariates, using K-means clustering did not change the final prediction model: the LGB covariate was also the only covariate in the final model. In the K-means method the number of cluster has to be fixed *a priori*. On the contrary, the number of clusters can be chosen *a posteriori* in the hierarchical clustering according to the results of the similarity, or according to one of the numerous existing criteria [**20**].

## Conclusion

Finally, one of the goals of this work was the use of the selected covariate in a usual model (e.g. linear regression). The results of such a model depend of course on the selection procedure. Thus, it is essential to use the less biased procedure such as this procedure based on the centroid criteria. Furthermore, when a cluster of covariates is summarized by a single covariate, amounts of information are discarded. Among all the covariates belonging to a cluster, the nearest covariate from the cluster centroid has to be selected to reduce amounts of discarded information. The maximum of the Pearson correlation sum is a useful criterion, and its practical use is very simple and fast. Thus, our method based on the centroid criterion is a simple and useful method to select covariates composing a set of predictors in the presence of missing data, in order to provide predictive model.

## Appendix 1: Determination of the centroid criteria

The previous statistical context depends on the following probabilistic model:

*X* = (*X_j_*), *j* = 1,…*p*

We note *ρ* _*jk*_ the correlation coefficient of Pearson between 2 variables, *X* _*j*_ and *X* _*k*_, *δ* ^2^ (*j, k*) the squared distance between the 2 variables *X* _*j*_ and *X* _*k*_, which is an Euclidean distance (and moreover a circum-Euclidean distance) [**15**]:

*δ* ^2^ (*j, k*) = (1– *ρ* _*jk*_), is a classical definition since the correlation matrix is positive semi-definite.

*δ* ^2^(*γ*.*j*)the squared distance between the variable *X* _*j*_ and the centroid *γ*.

For every Euclidean figure, the squared distance to the centroid is given by the Torgerson formula [**14**; **21**]:

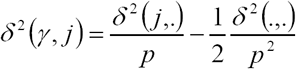

Where 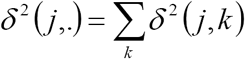 which depends only on *j*, and 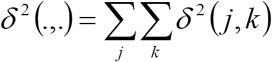 which is constant.

Now,

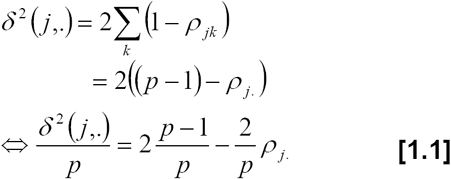

Where 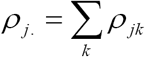

From [1.1] we can write that:

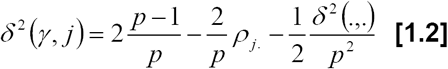

Now finding the nearest variable to the centroid is minimising *δ* ^2^(*γ*.*j*).

Therefore, from [1.2]:

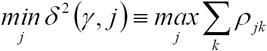

## Appendix 2: Almost-sure convergence

We consider the following statistical model:

Let *X* _*i*_, *i=1…n*, be a series of *n* i.i.d. random vectors, with a common continuous distribution, and admitting moments of second order (i.e. *E*[‖*X* _*i*_‖^2^ < ∞]).

Notations:

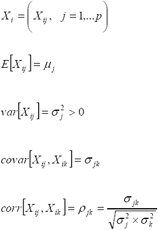

*δ* ^2^ (*j, k*) = (1– *ρ* _*jk*_) the squared distances from the theoretical model, which are circum-Euclidean distances

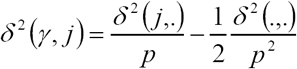 defining the centroid *γ*.

We can suppose the uniqueness of the nearest variable to the centroid:

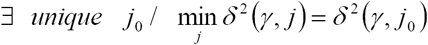

We now define the notations of the empirical model:

Empirical means *M*_*n,j*_

Empirical variances 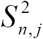

Empirical covariances *S*_*n, jk*_

Empirical correlation coefficients: *R*_*n, jk*_

Empirical squared distances: 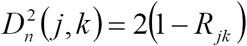

From the strong law of large numbers, we can note that:

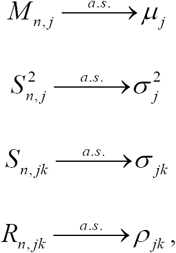

Since a continuous transformation preserves convergence, we can write that:

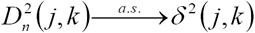

And

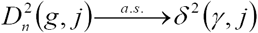

For a minimum and unique *δ* ^2^ (*γ, j*_*0*_), we have

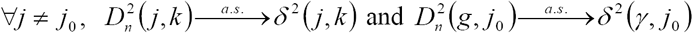

Let 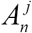 be the event 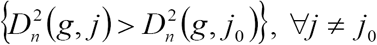,

### Proposition 1

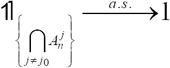

i.e. from a certain rank, the procedure built by the empirical statistics yields the right variable *j*_0_ with probability 1 (almost sure convergence).

### Demonstration

Let us start with a lemma:

### Lemma 1

Let *X* and *Y* be 2 random variables.

If 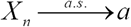 and 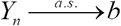, with *a<b*,

Then 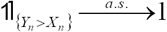

### Demonstration

If 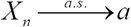 and 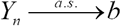, with *a<b*,

Then 

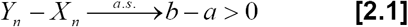

Now,

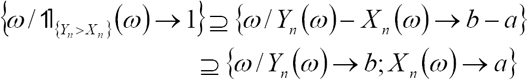

And from [2.1]

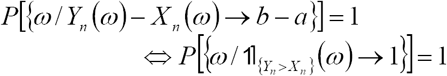

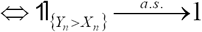

Then, from lemma 1:

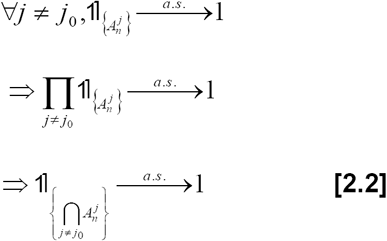

Moreover the almost sure convergence implies the convergence in probability.

Then,

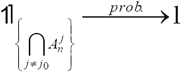

It is easy to see that this condition is equivalent to:

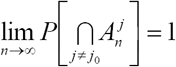

The probability, that 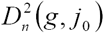 gives the minimum, tends to 1 when *n* is increasing, i.e. the probability to make the right choice tends to 1.

## List of abbreviations

CCSR: centroid method for covariate space size reduction;
CD: covariate deletion based on the sample size
CI95%: 95% confidence interval
Cst: model’s constant
DA: Data Augmentation algorithm
EM: Expectation-Maximization algorithm
MI: multiple imputations
p: p-value
R^²^: goodness of fit criterion: percent of variability explained by the model
SD: standard deviation
SI C-mean: simple conditional mean imputations (conditioned by Age), estimated from simple linear regression model, to missing data
SI UC-mean: simple unconditional mean imputations to missing data

**Biometric covariates**
ABO: angle formed by the straight line nasion-opisthion and the Francfort section
AFW: Weisbach angle
APA: alveolar profile angle
APN: nasal angle
ASP: nasio-sphenion-basion angle
ATO: foramen magnum angle
BEK: biorbital breadth
HMTG: mastoid height
HNZ: nasal height
JUB: bijugal breadth
LBH: basion-hormion length
LGB: glabella-basion length
LMA: maxillo-alveolar length
LNK: nasion-klition length
NKB: nasion-klition-basion angle
*ba*: basion
*gl*: glabella
*ns*: nasospinal
*pr*: prosthion

## Authors’ contributions

P Adalian carried out the anthropological data with the help of P Adalian.

J Gaudart run the statistical analysis, performed the simulations and the demonstrations, and wrote the initial draft.

G Leonetti initiated and supervised the anthropological study.

All authors read, corrected and approved the final manuscript.

## Acknowledgments

The authors gratefully acknowledge Bernard Fichet for his detailed and helpful comments, and Jean Giustiniani for helping carrying out the anthropological data.

## References

1. Hunsberger S, Murray D, Davis CE, Fabsitz RR: Imputation strategies for missing data in a school-based multi-centre study: the Pathways study. Stat Med 2001, 20:305–316.

2. Schafer JL, Olsen MK: Multiple imputations for multivariate missing-data problems: a data analyst’s perspective. Multivariate Behav Res 1998, 33(4):545–571.

3. Schafer JL: Multiple imputation: a primer. Stat Methods Med Res 1999, 8:3–15.

4. Clark TG, Altman DG: Developing a prognostic model in the presence of missing data: an ovarian cancer case study. J Clin Epidemiol 2003, 56:28–37.

5. Faris PD, Ghali WA, Brant R, Norris CM, Galbraith PD, Knudtson ML: Multiple imputation versus data enhancement for dealing with missing data in observational health care outcome analyses. J Clin Epidemiol 2002, 55:184–191.

6. Van Buuren S, Boshuizen HC, Knook DL: Multiple imputation of missing blood pressure covariates in survival analysis. Stat Med 1999, 18:681-694.

7. Joseph L, Bélisle P, Tamim H, Sampalis JS: Selection bias found in interpreting analyses with missing data for the prehospital index for trauma. J Clin Epidemiol 2004, 57:147–153.

8. Zhou XH, Eckert GJ, Tierney WM: Multiple imputation in public health research. Stat Med 2001, 20:1541–1549.

9. Schafer JL: Analysis of incomplete multivariate data. Edited by Cox DR, Isham V, Keiding N, Reid N, Tong H. Boca Raton: Chapman and Hall; 1999. [Monographs on Statistics and Applied Probability, vol 72.]

10. Adalian P, Piercecchi-Marti MD, Boulière-Najean B, Panuel M, Leonetti G, Dutour O: Nouvelle formule de détermination de l’âge d’un fœtus. C R Biol 2002, 325: 261–269.

11. Streeter GL: Weight, sitting height, head size, foot length and menstrual age of the human. Contrib Embryo Carnegie Inst 1920, 11:143.

12. Giustiniani J: Ontogénèse crânio-faciale et ses applications en anthropologie médico-légale. PhD thesis, Aix-Marseille university, Anthropology and Forensic Sciences Unit; 2005.

13. Martin R, Saller K: Lehrbuch der Anthropologie 1/2. Stuttgart: G Fisher Press; 1957.

14. Torgerson WS: Theory and methods of scaling. New York: Wiley Press; 1958.

15. Fichet B: Distances and Euclidean distances for presence-absence characters and their application to factor analysis. In: Multidimensional data analysis. Edited by De Leeuw J, Heiser W, Meulman J, Critchley F. Leiden: DSWO Press; 1986;23–46.

16. Fisher RA: Frequency distribution of the values of the correlation coefficient in samples from an indefinitely large population. Biometrika 1915, 10:507–521.

17. Soper HE, Young AW, Cave BM, Lee A, Pearson K: On the distribution of the correlation coefficient in small samples. Biometrika 1916, 11:328–413.

18. Olivier G, Demoulin F: Pratique anthropologique à l’usage des étudiants. Paris: Université Paris VII; 1976.

19. Collins LM, Schaffer JL, Kam CH: A comparison of inclusive and restrictive strategies in modern missing data procedure. Psychol methods 2001, 6(4):330–351.

20. Roux M: Algorithmes de classification. Paris: Masson; 1985.

21. Van der Waerden BL: Statistique mathématique. Paris: Dunod Press; 1967.

